# The genetic basis of adaptation through the evolution of self-fertilization

**DOI:** 10.1101/2021.02.11.430833

**Authors:** Kuangyi Xu

## Abstract

Although adaptation can be realized through the fixation of beneficial alleles that increase viability, many plant populations may adapt through the evolution of self-fertilization, especially when pollination becomes inefficient. However, the genetic basis of adaptation through the evolution of selfing remains unclear. Using population genetic models, I study adaptation through the fixation of alleles that increase the selfing rate (selfing modifiers) from new mutations or/and standing variation. For adaptive alleles unrelated to selfing, it is known that selfing promotes adaptation from a new mutation only when the beneficial alleles are recessive, and the probability of adaptation from standing variation is nearly independent of dominance, and always decreases with the selfing rate. In contrast, for adaptation through the evolution of selfing, when it occurs by fixation of a newly arisen mutation, a population that already has a high selfing rate may be more (less) likely to adapt than outcrossers even when the modifier is dominant (recessive) if the modifier is weakly (strongly) selected. Also, adaptation from standing variation is more likely through recessive modifier alleles, with the highest fixation probability found in partially selfing populations, but fixation is fastest when dominance is intermediate. When there are multiple modifiers, adaptation through new mutations is more likely when selfing is controlled by few large-effect rather than many slight-effect modifiers. This study suggests that to understand the genetic basis of adaptation, it is necessary to determine the ecological and genetic advantages of adaptive alleles.

**Significance statement:** This study, by deriving the selective coefficient and effective population size, investigated the genetic basis of adaptation through fixation of modifier alleles that increase the selfing rate, which is shown to differ in several aspects from that through evolution of mating-unrelated alleles. Specifically, when adaptation is from new mutations, the dominance of a selfing modifier allele below which selfing increases the fixation probability depends on the strength of pollen limitation and pollen discounting. Adaptation from standing variation is more likely through recessive modifier alleles and in populations with an intermediate selfing rate. This work suggests it is important to have a mechanistic understanding of how adaptive alleles increase individual fitness in environment that is unfavorable to the population.

## Introduction

Adaptation often involves fixation of alleles that increase individual fitness, either through new beneficial mutations, or through alleles that are previously present in the population before an environmental change, often called standing genetic variation (Hermisson and Pennings 2005, Barret and Schluter 2008, Orr 2010, Lai 2019). In many plant populations, adaptation may occur through the evolution of their mating systems, especially when environmental change reduces mating success (Barret et al. 2008, Eckert et al. 2010, Pannell 2015, Cheptou 2019). Although our understanding of adaptation through mating-unrelated beneficial alleles has been greatly promoted by previous studies (Orr 2001, Hermisson and Penings 2005, Barret and Schluter 2008, Orr 2010, Ronfort and Glemin 2013, Glemin and Ronfort 2013), how the genetic properties of alleles which modify the selfing rate (hereafter referred as selfing modifiers or modifiers) influence the probability of successful adaptation remains unclear.

Adaptation through the evolution of mating systems may be particularly common for today’s plant populations, since an increased level of human disturbance and widespread declines in pollinator populations have intensified limitation of pollen (Larson and Barrett 2000, Knight et al. 2005) and reduced plant reproductive success (Eckert et al. 2010, Thomann et al. 2013). Compared to outcrossing, selfing depends less on extrinsic factors, and thus may offer assurance for seed reproduction when pollination service is scarce (Holsinger 2000, Eckert et al. 2006). Indeed, an increased level of self-fertilization has been widely documented in response to various environmental conditions, including human disturbances (Eckert et al. 2010), habitat fragmentation (Aguilar et al. 2006), range expansion (Levin 2012), and colonization (Barrett et al. 2008, Pannell 2015). Also, the evolution of selfing is often associated with morphological changes in several floral characters (Goodwillie et al. 2010, Sicard and Lenhard 2011), which are analyzed to be controlled by multiple loci with both large and small effects (Fishman et al 2002, 2015, Goodwillie et al. 2006).

Adaptation through the fixation of selfing modifiers differs from that of mating-unrelated alleles in several aspects. First, the selective advantage of a modifier allele depends on both genetic and ecological factors. Regarding genetic factors, selfing enjoys a transmission advantage over outcrossing (Fisher 1941), but suffers the cost of inbreeding depression (Charlesworth and Willis 2009). In the ecological aspect, selfing imposes a cost as it may reduce opportunities for pollen export, often called pollen discounting (Harder and Wilson 1998, Porcher and Lande 2005), but is more advantageous when pollen limitation becomes severe. Second, in a finite population, a selfing modifier should have a different effective population size from a mating-unrelated locus, since the selfing rate is polymorphic at the modifier locus, and during the fixation process, the level of inbreeding is no longer constant but changes with the frequency of the modifier. Third, although previous studies show that the fixation probability of mating-unrelated alleles will often be lower in populations with higher selfing rates (Caballero and Hills 1992, Glemin and Ronfort 2013), this may not hold for selfing modifier alleles, as the selective advantage and effective population size of a modifier may depend on the selfing rate that is already in the population The answer to this question may also help us understand the speed of the evolution of selfing rates during the transition from outcrossing to selfing, which is commonly seen in plant populations (Igic and Busch 2013).

To examine the genetic basis of adaptation through the evolution of selfing, I first derive the effective selection coefficient and the effective population size for a selfing modifier. I then use the diffusion approximation to obtain the fixation probability of such a modifier from a new mutation and from standing variation. I consider adaptation through fixation at a single modifier locus in three demographic situations: a constant population, a declining population, and a population after a bottleneck. For adaptive alleles unrelated to selfing, selfing promotes adaptation from a new mutation only when it is recessive, and the probability of adaptation from standing variation is independent of dominance. In contrast, for adaptation through the evolution of selfing, when adaptation is from a new mutation, selfing may inhibit adaptation even when the modifier is recessive if it is strongly favored, and can facilitate adaptation for dominant, weakly-favored modifiers, while adaptation from standing variation is more likely through recessive modifiers. Generally, partial-selfing populations tend to have the highest fixation probability of selfing modifiers, suggesting a nonlinear evolutionary rate from outcrossing to selfing. An extension to multiple loci suggests that adaptation through the evolution of selfing is more likely when selfing is controlled by few large-effect modifiers rather than many small-effect modifiers.

## Models and results

I consider a selfing rate modifier locus with two alleles *A* and *a* in a hermaphroditic, diploid population with non-overlapping generations. Asexual reproduction, that is, apomixis (Asker and Jerling 1992), which only happens in less than 1% of the species (Mogie 1992), is not considered and thus seeds will be produced from either selfing or outcrossing. The population already has a background selfing rate *r*_0_, and the selfing rate of the three genotypes *AA, Aa* and *aa* are *r*_0_ + *r, r*_0_ + *hr* and *r*_0_, where *h* is the dominance coefficient. It is assumed that the genetic properties of the selfing modifier locus are independent of the environment, so *r* and *h* are the same before and after pollen limitation. The inbreeding depression in the population is *d* and the level of pollen discounting is *c*, which means for an individual with a selfing rate *r*_*i*_, a proportion *r*_*i*_*c* of its pollen will not be disseminated compared to a completely outcrossing individual. The population suffers from pollen limitation with a strength *m*, i.e., a proportion *m* of its ovules which should have been outcrossed will not be fertilized. Since the number of viable offspring from selfing relative to outcrossing is (1 − *d*)/(1 − *m*), I can define an effective inbreeding depression, which incorporates the effect of pollen limitation as

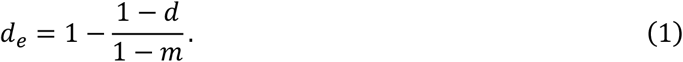

Evolution towards a higher selfing rate will increase the average fitness only when 1 − *m* < 1 − *d*, so that *d*_*e*_ < 0, but *d*_*e*_ > 0 will also be considered for a thorough analysis.

To model drift at a modifier locus, the key is to obtain the effective selective coefficient and the effective population size, based on which the fixation probability of alleles starting from any frequency can be calculated. This is done by first considering the frequency change and its variance after one discrete generation (see section 1 in SI), and then using the diffusion approximation by considering an infinitesimal time interval *dt* → 0. The overall allele frequency change is obtained by a weighted average of changes among selfed and outcrossed seeds. For outcrossed seeds, the allele frequency is the average of those in the pollen and ovule pool. The overall variance is the sum of two components: variance in selfed and outcrossed offspring given that the proportion of selfed seeds is fixed, and variation of the actual proportion of selfed offspring. The first component is calculated by assuming the number of selfed and outcrossed offspring per individual is subject to a Poisson distribution. After that, the fixation probability of a modifier allele starting from any frequency can be obtained. The expected fixation probability from standing variation is integrated over the initial frequency distribution under the selection-drift-mutation equilibrium before pollen limitation. For declining populations, the fixation probability is modified to account for the reduction of effective population size caused by demographic decline, and the population goes to extinction when all pre-existing alleles and all mutations that appears before the population size declines to 0 will not go fixation. I also extend the model to investigate adaptation after a bottleneck, the effect of genetic architecture and coevolution of inbreeding depression. A full presentation can be found in the Methods section.

In the section 2 of SI, I show that the effective selection coefficient *R* the effective population size of the selfing enhancer allele *A* are

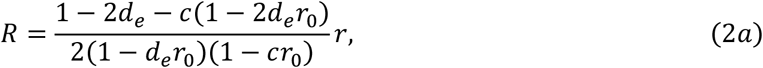

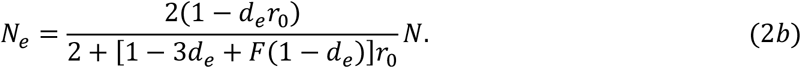

*F* is the inbreeding coefficient, which measures the deviation of the frequency of homozygotes from the Hardy-Weinberg equilibrium. Although *F* will change with time, it approaches the equilibrium much more quickly than the allele frequency changes (Hartfield and Glemin 2016). Therefore, *F* is first approximated with its equilibrium value, which depends on the allele frequency. However, it is shown that a secondary approximation of *F* as a frequency-independent constant works well (Table S1), given by

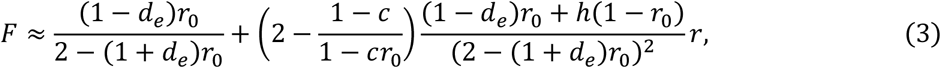

Equation (2a) shows the selective advantage depends on the background selfing rate, and unlike mating-unrelated loci, equations (2b) and (3) show that the effective population size depends on the effect of the modifier *r. N*_*e*_ also decreases with *d*_*e*_ and *c* (see section 5 in SI), and thus is regulated by ecological factors. In contrast, the effective population size of mating-unrelated alleles should increase with *d*_*e*_ since *d*_*e*_ lowers the level of inbreeding (Caballero and Hill 1992).

### Adaptation in a constant population

When fixation is from a single new modifier mutant, equation (7) in the Methods section shows that fixation is more likely through dominant modifiers. However, whether a higher background selfing rate will promote or inhibit fixation depends on the dominance coefficient. For a selfing modifier, the critical dominance coefficient *h*_*c*_ below which selfing increases the fixation probability (obtained from equation (8) in Methods) is

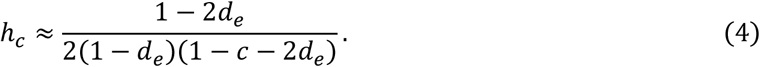

As shown in Fig. 1, *h*_*c*_ is higher than 0.5 under weak pollen limitation and strong pollen discounting (large *d*_*e*_ and *c*). The critical dominance above which adaptation through evolution of selfing is less likely in selfers than outcrossers becomes more restricted compared to adaptation through mating-unrelated alleles. On the contrary, when the modifier is strongly favored (*d*_*e*_ and *c* small), *h*_*c*_ is often smaller than 0.5. Therefore, when the evolution of selfing will increase the average fitness (i.e., *d*_*e*_ < 0), outcrossers may be much more likely to adapt than selfers. In contrast, *h*_*c*_ is always 0.5 for mating-unrelated alleles. This difference is mainly because the selective advantage of a selfing modifier depends on the background selfing rate (see equations (2) and (8) Methods).

**Figure 1.**
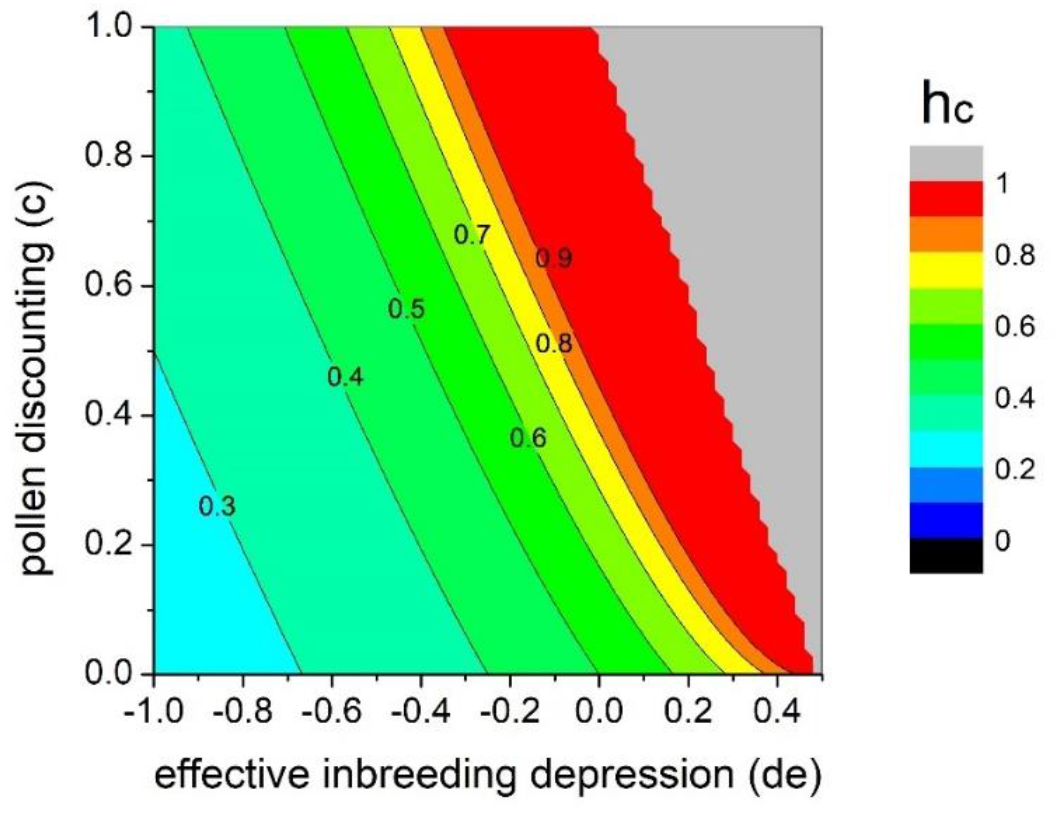
Effects of pollen discounting *c* and effective inbreeding depression *d*_*e*_ on the value of critical dominance *h*_*c*_. Results are calculated based on equation (4), but the value is set to be 1 if *h*_*c*_ > 1. The grey area is where for all levels of dominance, the selfing modifier is selected against in a completely outcrossing population and thus cannot fix, but favored in a highly selfing population (see equation (8)).

When fixation is through standing variation, for mating-unrelated alleles, the fixation probability is nearly independent of the dominance (Orr and Betancourt 2001), which holds for both outcrossing and selfing populations (Glemin and Ronfort 2013). However, for selfing modifier alleles, fixation from standing variation is more likely through recessive alleles, as shown in Fig. 2. This pattern is more prominent when the background selfing rate *r*_0_ is intermediate, when the modifier is more strongly selected against before pollen limitation, and when the population size is larger (see Fig. S1). Also, for mating-unrelated alleles, the fixation probability from standing variation decreases with the selfing rate (Glemin and Ronfort 2013). For adaptation from the evolution of selfing, this result usually holds when the modifier is not very recessive (*h* = 0.5, 0.9 in Fig. 2), but when the modifier allele is recessive enough, the fixation probability will be highest when the background selfing rate is at an intermediate level (*h* = 0.1 in Fig. 2). However, when pollen limitation is weak and pollen discounting is strong (Fig. 2(c)), in which case selfing is more likely to enhance the fixation probability after pollen limitation (see Fig.1), fixation probabilities increase almost monotonically with the background selfing rate. The above findings are robust to the effect of the selfing modifier *r*, as the fixation probability is nearly independent of *r* (Fig. S2).

**Figure 2.**
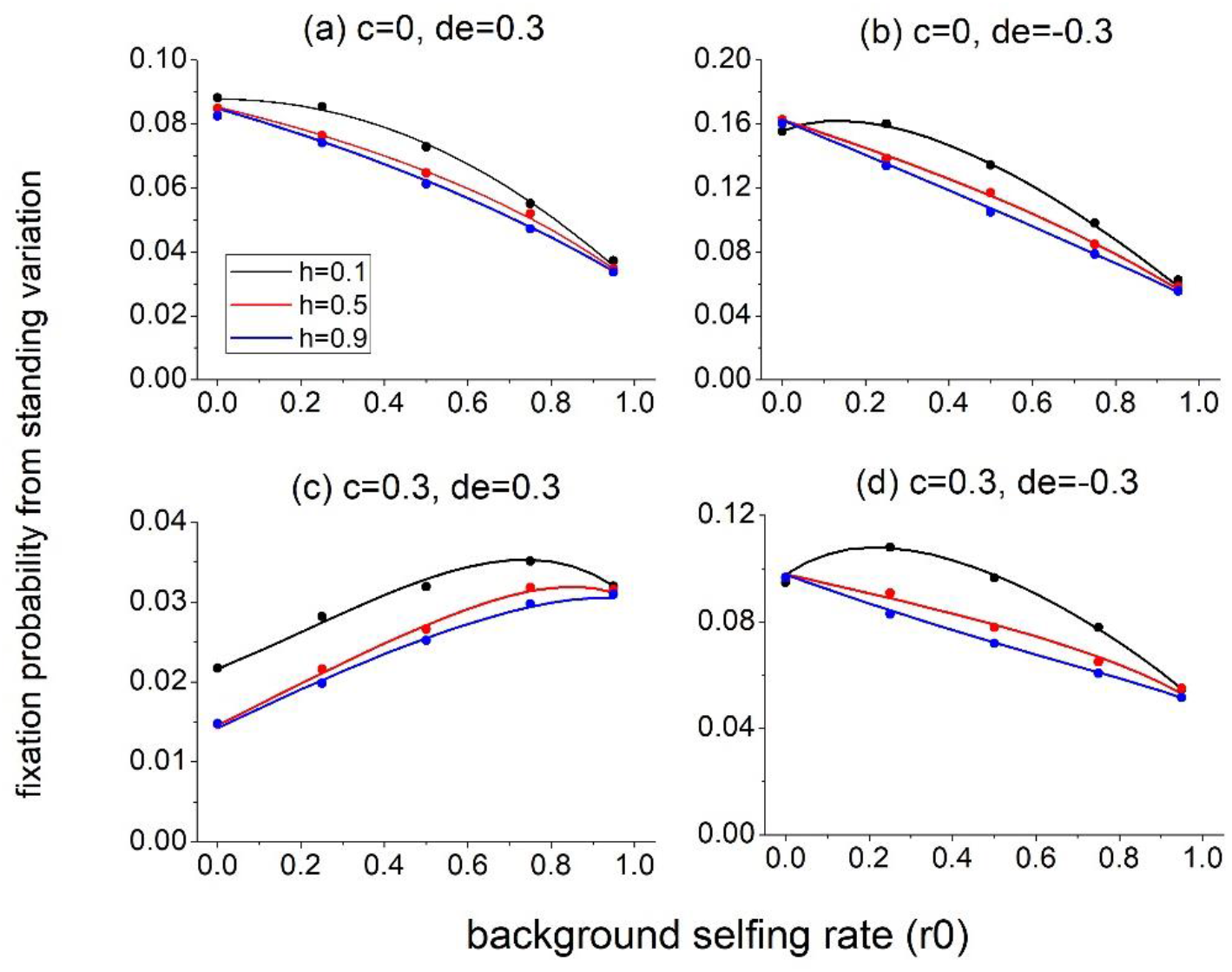
Effects of background selfing rate on fixation probability from standing variation in a constant population. Lines are model predictions from equation (10), and dots are simulation results. For all panels, *r* = 0.01, *d* = 0.6, *N* = 20000, *v*_1_ = *v*_2_ = 10^−6^.

Despite a higher fixation probability for recessive modifier alleles when adaptation is from standing variation, generally, fixation is fastest when the allele has intermediate dominance, while it is slowest for dominant alleles (Fig. S3). However, when the allele is under weak selection fixation is quickest for recessive alleles (Fig. S3(c)). The above findings are robust to *r* (Fig. S4). As a contrast, for mating-unrelated alleles, more dominant alleles will go to fixation more quickly (Glemin and Ronfort 2013).

The above differences mainly result from the fact that the evolution at a selfing modifier locus will change the level of inbreeding at the locus itself. At the selection-mutation-drift equilibrium before pollen limitation, because selfing enhancer alleles are exposed more in homozygotes, they will be kept at an even lower frequency than mating-unrelated alleles of the same selective disadvantage. Moreover, a higher dominance will strengthen the selection in two ways. First, it directly increases the selfing rate of the heterozygote. Second, an enhanced selfing rate of the heterozygotes further increases the homozygosity, which indirectly increases the strength of selection. Therefore, compared to dominant mating-unrelated alleles, which increases strength of selection only through heterozygotes, a dominant selfing modifier will be more strongly selected against before pollen limitation, and although it will also be more strongly favored after pollen limitation, this advantage usually cannot compensate for the disadvantage before pollen limitation.

### Adaptation in a declining population

The previous section considers populations that are still able to persist and maintain a constant size after pollen limitation. However, some populations may not produce sufficient offspring and thus will suffer a demographic decline and go extinct. To investigate adaptation in declining populations, I consider a population which has an expected offspring number 1 − *λ* after pollen limitation, and will be successfully rescued when the selfing enhancer allele *A* goes to fixation.

For the case of evolutionary rescue occuring through new mutations, equation (16) in the Methods shows that the critical dominance at which selfers and outcrossers have the same rescue probability is higher than that in a constant population. Therefore, demographic decline favors selfers to be more likely to adapt than outcrossers.

For rescue from standing variation, demographic decline favors adaptation through dominant alleles compared to a case with the same conditions in a constant population. This can be seen, for example, by comparing *r*_0_ = 0 in Figs. 3(a) and 2(a). This difference is more prominent when *r* is small (Fig. S5). Moreover, for all levels of dominance (Fig. 3(b)), the proportion of rescue from standing variation is lower for dominant alleles, and decreases with the background selfing rate, suggesting an increased relative contribution from new mutations in selfing populations. However, how dominance influences the overall rescue probability from both sources should depend on the relative contribution from standing variation versus new mutations. Rescue will be more likely through recessive alleles when the contribution from standing variation is larger, which happens under a smaller effect *r* (Fig. S5), a larger initial population size and a faster decline rate (Glemin and Ronfort 2013).

**Figure 3.**
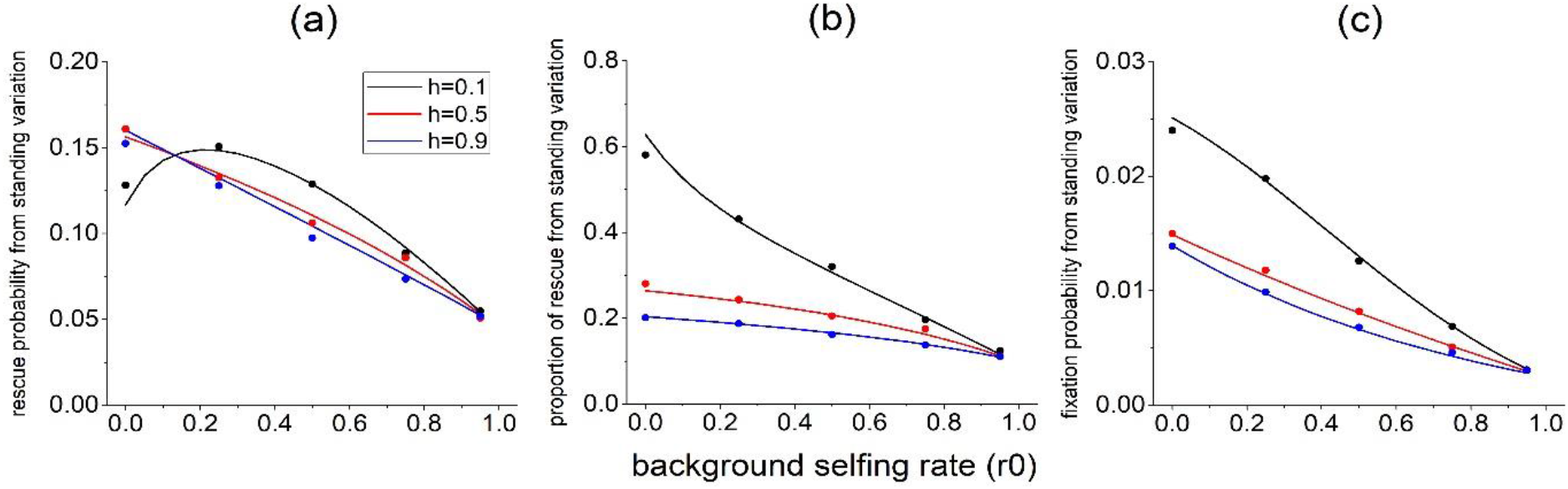
Effects of the background selfing rate on the rescue probability in declining populations (panels (a), (b)) and populations suffering a bottleneck (panel (c)). Lines in panels (a), (b) and (c) are model predictions from equation (14), (17) and (18), respectively, and dots are simulation results. For panels (a) and (b), *r* = 0.05, *d* = 0.6, *d*_*e*_ = −0.3, *c* = 0, *N*_0_ = 20000, *λ* = 0.002, *v*_1_ = *v*_2_ = 10^−6^. For panel (c), *r* = 0.01, *d* = 0.6, *d*_*e*_ = −0.3, *N*_0_ = 20000, *N*_*c*_ = 500, *v*_1_ = *v*_2_ = 10^−6^.

### Adaptation after a bottleneck

In the previous sections, I consider that the pollen limitation happens in the original habitat. However, the evolution of self-fertilization is often associated with bottleneck effects to some degree (Eckert et al. 2010, Aguilar et al. 2006, Levin 2012, Barrett et al. 2008, Pannell 2015). I consider a population of an ancestral size *N*_0_ that suffers from a bottleneck effect of *N*_*c*_ individuals in a new environment. equation (7) indicates that a bottleneck will have little effect on the fixation probability of a new mutant. For adaptation from standing variation, compared to a constant population, a bottleneck lowers the fixation probability (compare the y axis in Figs. 3(c) and 2(a)). Fixation is even more biased for recessive alleles, especially when the background selfing is low, which can be easily seen by comparing Fig. 3(c) with Fig. 2(a) at *r*_0_ = 0. In contrast, the fixation probability is usually insensitive to dominance for mating-unrelated alleles (Fig. S6). Moreover, for all levels of dominance, the fixation probability always decreases with the background selfing rate. It is often considered that self-compatible populations are more likely to establish after long-distance dispersal, known as Baker’s law (Baker 1951, Cheptou 2012). However, it is often misinterpreted as an association between higher selfing rates and successful colonization. Our result indicates that establishment of a colony through the genetic evolution of selfing is more probable for outcrossing populations, which supports the view of Pannell et al. (2015) that Baker’s law should be interpreted in terms of an enhanced capacity of selfing associated with colonization.

### Effects of the genetic architecture of selfing on adaptation

As selfing is often controlled by multiple modifier loci, it is critical to ask whether adaptation is more likely when the selfing rate is controlled by many slight-effect modifier loci or few large effect loci. For adaptation from standing variation, as the fixation probability at a single locus is insensitive to its effect *r* (Fig. S2), adaptation is more likely when there is a larger number of loci. However, the answer is not clear for adaptation from new mutations. Although genetic architecture of many slight-effect loci has a higher overall mutation rate, the fixation probability of a single modifier is lower. Here I assume modifier loci act additively on the selfing rate, as selfing rates sometimes have an approximately linear relationship with phenotypic values of several quantitative characters (Karron et al. 1997, Strand et al. 2017, Fishman et al. 2002, 2015). equation (19) in Methods and Fig. S7 show that in constant populations, adaptation from new mutations is always faster under few large-effect modifier loci, consistent with the empirical findings (Bodbyl Roels and Kelly 2011), and rescue of declining populations is also more likely through the genetic architecture of few large-effect loci (see discussion below equation (19)).

### Incorporating the genetic basis of inbreeding depression

Up to now, inbreeding depression is treated as a constant parameter. However, inbreeding depression is caused by deleterious mutations and will coevolve with the mating system. Previous models and empirical evidence have shown that a higher selfing rate will lower the inbreeding depression at the equilibrium (Lande and Schemske 1985, Roze 2015). Moreover, selection against deleterious mutations may also inhibit fixation of a linked beneficial mutation, known as background selection (Charlesworth et al. 1993, Charlesworth 2012), which is more severe in selfing populations due to a reduced effective recombination rate.

When inbreeding depression is fixed, adaptation probability tends to either decreases or increases monotonically with the selfing rate. In contrast, under the coevolution of inbreeding depression, Fig. 4 shows that for both adaptation from new mutations and standing variation, the fixation probability is highest at an intermediate selfing rate, suggesting that a trade-off between the benefit of lower inbreeding depression and the cost of higher background selection. More specifically, the selfing rate corresponding to the highest fixation probability is higher for recessive alleles when fixation is from a new mutation (Fig. 4(a)), but lower when it is from standing variation (Fig. 4(b)). Although the rate of the transition from outcrossing to selfing has been estimated at the phylogenetic scale (Igic and Busch 2013), the above result implies that at a smaller scale during this transition, the evolution of self-fertilization accelerates when the population initially starts from being predominantly outcrossing, while slows down later when it approaches predominantly selfing.

**Figure 4.**
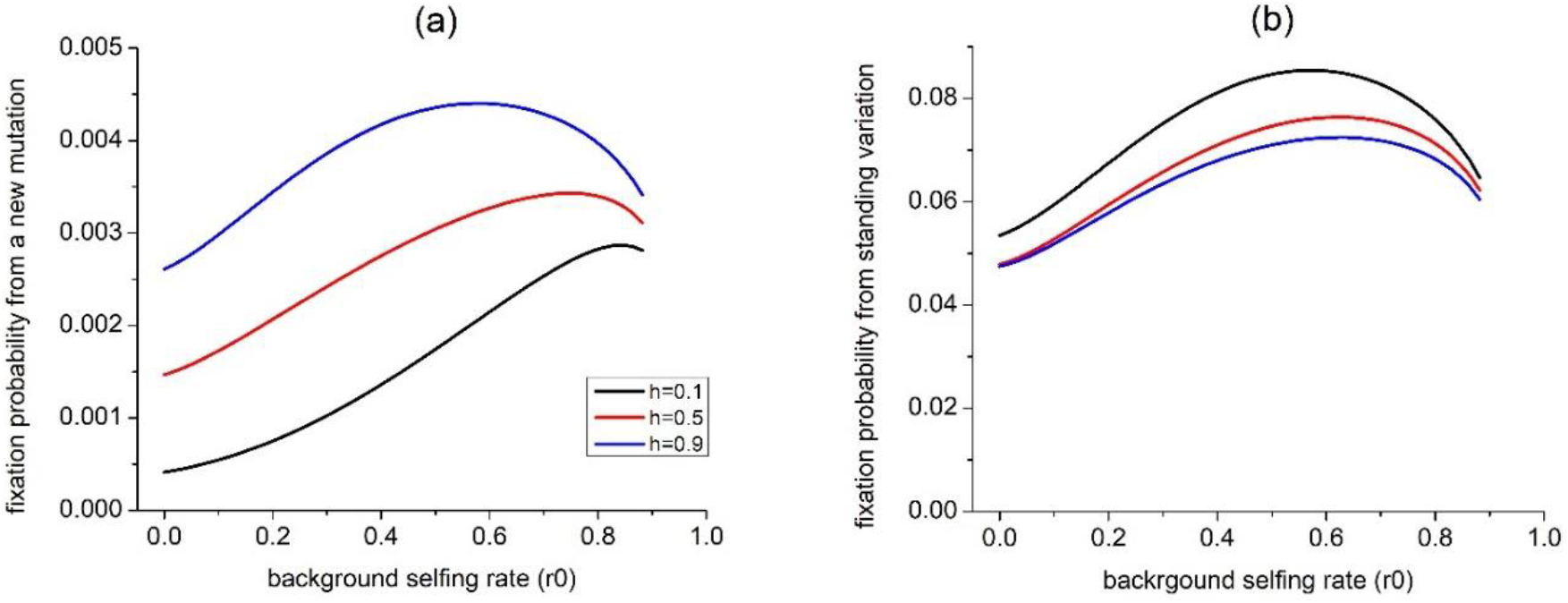
Effects of the background selfing rate on the population survival probability from a new mutation (panel (a)) and standing variations (panel (b)) when incorporating the genetic basis of inbreeding depression and background selection. Parameters related to deleterious mutations are *U* = 0.4, *h*_*d*_ = 0.2, *s* = 0.1, *d*_0_ = 0.4, *L* = 10; and other parameters are *r* = 0.01, *c* = 0, *m* = 0.5, *N* = 20000, *v*_1_ = *v*_2_ = 10^−6^.

**Table 1.** Comparison of the genetics of adaptation for mating-unrelated alleles with selfing modifier alleles. *h*_*c*_ is the critical dominance above which fixation probability is lower in selfers than outcrossers.

|  | Mating-unrelated allele | Selfing modifier allele |
| --- | --- | --- |
| Adaptation from new mutations in a constant population | 1. Fixation probability is higher for dominant alleles.<br>2. $h_c = 0.5$ | 1. Fixation probability is higher for dominant alleles.<br>2. $h_c$ can vary from 0 to 1, depending on the strength of pollen limitation and pollen discounting. |
| Adaptation from standing variation in a constant population | 1. Fixation probability is nearly independent of dominance.<br>2. Fixation probability decreases with the selfing rate.<br>3. Fixation is quickest for dominant alleles. | 1. Fixation probability is higher for recessive alleles.<br>2. Fixation probability is highest at an intermediate selfing rate.<br>3. Fixation is quickest when alleles are intermediate dominant, while slowest for dominant alleles; |
| Adaptation in declining populations | 1. $h_c$ increases.<br>2. Rescue is more likely through dominant alleles.<br>3. Rescue probability from standing variation is lower in selfers | 1. $h_c$ increases.<br>2. Rescue is more likely through dominant alleles.<br>3. Rescue probability from standing variation is lower in selfers. |
| Adaptation in populations after bottleneck | 1. Fixation probability is insensitive to dominance.<br>2. Adaptation is less likely in selfers. | 1. Recessive alleles have a much higher fixation probability.<br>2. Adaptation is less likely in selfers. |
| Genetic architecture of adaptation | N/A | Adaptation from standing variation more likely when selfing is controlled by many small-effect loci, while adaptation from new mutations should be faster and more likely through few large-effect loci. |
| Under the coevolution of inbreeding depression | N/A | The fixation probability from both new mutations and standing variation is highest at an intermediate selfing rate. |

## Conclusion

While the genetic basis of adaptation through the fixation of mating-unrelated adaptive alleles has been well studied (Haldane 193, Orr 2001, Hermisson and Pennings 2005, Glemin and Ronfort 2013), it is not clear whether these results can apply to the case when adaptation is through the evolution of sefl-fertilization. In this study, I investigate fixation of selfing modifier alleles from new mutations and standing variation in populations under different demographic situations. Although for mating-unrelated alleles, selfing increases adaptation from new mutations as long as they are recessive, selfing can lower the fixation probability of a recessive modifier if pollen limitation is strong, but makes dominant modifier more likely to fix under weak pollen limitation (Fig. 1). On the other hand, adaptation from standing variation is more likely through recessive modifier alleles (Fig. 1), which seems to be supported by previous genetic analysis of the rapid evolution of selfing rates in *Mimulus guttatus* (Bodbyl Roels and Kelly 2011). However, fixation is fastest for intermediate dominant modifier alleles (Fig. 2). Generally, adaptation is more likely through dominant alleles under a demographic decline, consistent with adaptation while a bottleneck effect favors recessive alleles.

These characteristics of adaptation through the evolution of mating systems result from two factors. First, a selfing modifier allele has a different inbreeding coefficient from mating-unrelated alleles, as it directly influences the level of inbreeding at the locus itself. Second, dominance has two-fold effects, as it directly increases the selfing rate of the heterozygote, which further increases the homozygosity. Although the dominance coefficient distribution of mating-unrelated alleles is generally unknown (Orr 2010), for selfing modifier alleles, it may be possible to estimate dominance by combining QTL analyses of selfing-related characters (Fishman et al. 2002, 2014) with studies on the association between selfing rate and phenotypic values. This work also provides insights on how the selfing rate of the population affects adaptation through evolution towards a higher selfing rate. Generally, a higher selfing rate limits the further increase of self-fertilization, but this may be counterbalanced by the coevolution of deleterious mutations in the genome (Fig.5).

Although I contrast adaptation through the evolution of mating system with that through increase of viability or fertility, adaptation may involve both paths through fixation of alleles that simultaneously influence several aspects of individual fitness. For example, an allele that reduces the flower display may not only allow more resources to be allocated for fertility (Sargent et al. 2007), but also increase the selfing rate (Sicard and Lenhard 2011). On the other hand, alleles that increase selfing may impose a fitness cost. For example, reduction of flower size may be caused by deleterious mutations that inhibit development (Kelly and Willis 2001). This pleiotropy may make the genetic basis underlying adaptation more complicated. All in all, this work suggests the importance of a mechanistic understanding of the source of the selective advantage of adaptive alleles.

## Methods

### Adaptation in a constant population size

The fixation probability of a selfing modifier allele with an initial frequency *x* is given by (Kimura 1962)

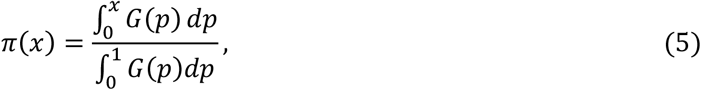

where

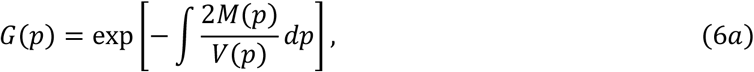

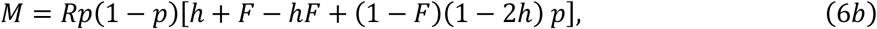

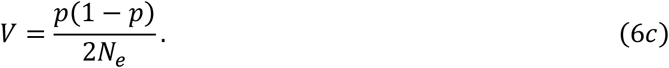

*M* and *V* are the expected change of the allele frequency and its variance under the limit of an infinitesimal time interval. When *N*_*e*_ (*h* + *F* − *hF*)*R* ≫ 1, the fixation probability can be approximated as

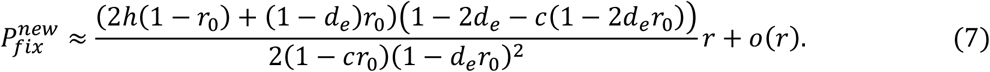

The fixation probability of a selfing modifier mutant in a completely outcrossing (*r*_0_ = 0) and a nearly completely selfing (*r*_0_ = 1 − *r*) population can be approximated as

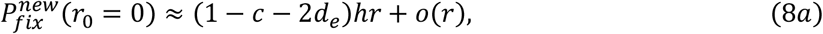

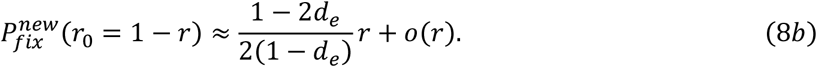

When the above two equations are equal, we can solve for the critical dominance *h*_*c*_ at which the fixation probability is the same for outcrossers and selfers, given in equation (4).

For adaptation from standing variation, denote the mutation rate from allele *a* to *A* as *v*_1_, and the backward mutation rate as *v*_2_. Before pollen limitation happens, allele *A* is selected against, and in a finite population under selection-drift-mutation balance, the stationary distribution of frequency of allele *A* is (Ewens 2012)

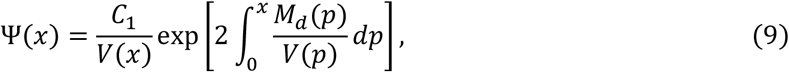

where *C*_1_ is a constant which normalizes the integral as 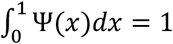, and *M*_*d*_(*p*) = *M*(*p*) + (1 − *p*)*v*_1_ − *pv*_2_, with *d*_*e*_ replaced by *d*. Therefore, the expected fixation probability from standing genetic variation is (Orr 2001)

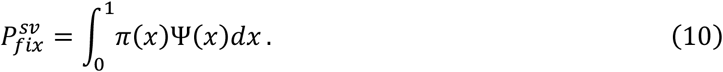

### Adaptation in a declining population

Suppose the maximum number of offspring an individual can produce is *W*_0_, the absolute fitness of *aa* individuals after pollen limitation happens is

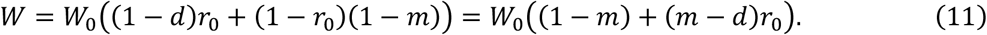

The absolute fitness of *AA* individuals is

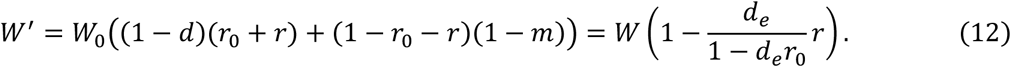

Therefore, for the evolution of selfing to rescue the population (*W*′ > 1), it requires *r* > (1 − *d*_*e*_*r*_0_)(1 − *W*^−1^)/*d*_*e*_. Denote the rate of demographic decline as *λ* = 1 − *W*, the fixation probability from a single mutant is reduced to (Glemin and Ronfort 2013)

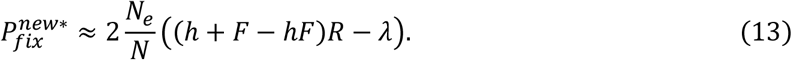

We assume the strength of pollen limitation is independent of the population size, so the probability for a population starting from an initial size *N*_0_ to be rescued by new mutations is approximately (Orr and Unckless 2008)

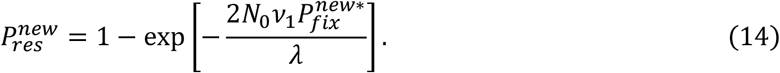

To obtain the critical dominance at which the fixation probability is the same for outcrossers and selfers, note that the rescue probability from new mutations in a completely outcrossing (*r*_0_ = 0) and a nearly completely selfing (*r*_0_ = 1 − *r*) population are

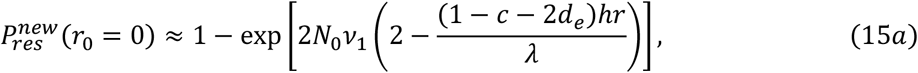

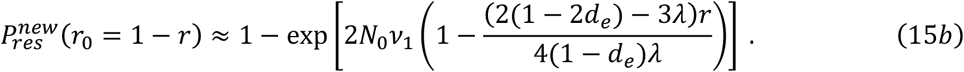

Therefore, the critical dominance is

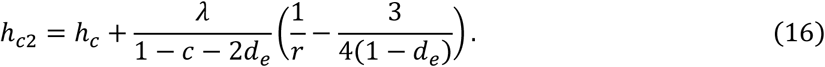

Since *r* is assumed to be small and *d*_*e*_ < 0.5, the second term is usually positive, and thus *h*_*c*2_ > *h*_*c*_. The probability for the population to be rescued by standing variation is approximately

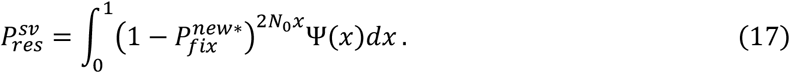

The population will go to extinction when alleles from both standing variation and new mutations cannot fix, so the overall rescue probability is 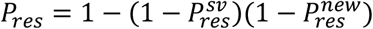, and the relative contribution to rescue from standing variation is 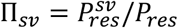.

### Adaptation after a bottleneck

Here we consider a population with an initial size *N*_0_ that suffers a bottleneck, so that the population size is suddenly reduced to *N*_*c*_. Suppose the allele frequency in the ancestral population is *x*, given that *N*_0_ ≫ *N*_*c*_, the probability that the allele frequency after the bottleneck *x*′ is approximately a binomial distribution. Therefore, the expected fixation probability from standing variation is approximately

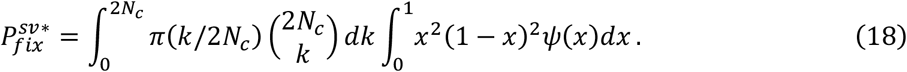

### Adaptation from multiple loci

For a certain overall increase of the selfing rate Δ*r*, I suppose there are *k* identical modifier loci, so the effect of each modifier is *r* = Δ*r*/*n* when modifiers act additively. For adaptation from new mutations, if the waiting time for a mutation to happen is longer than the time for a modifier to sweep to high frequency, i.e., 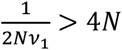. In this case, modifiers fix successively, and the time for all the modifiers to fix is

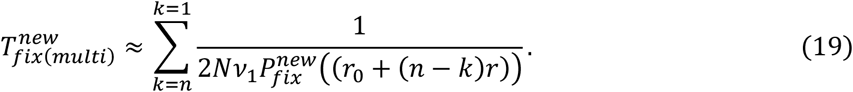

Fig. S7 shows that 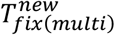 increases with *n*. For rescue of a declining population, if fixation of a single modifier is enough to stop demographic decline, the probability of rescue from multiple loci is still given by equation (14), with *v*_1_ replaced by *nv*_1_. Therefore, the effect of genetic architecture on rescue can be assessed from 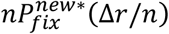, which decreases with *n*.

### Incorporating the genetic basis of inbreeding depression

It has been firmly established that inbreeding depression is mainly contributed to by two categories of deleterious mutations (Charlesworth and Willis 2009): slightly deleterious, partially dominant mutations, and large-effect, highly recessive lethal and sublethal mutations. Slightly deleterious, partially dominant mutations also cause background selection (Charlesworth 2012). I denote the inbreeding depression contributed by highly recessive large-effect mutations as *d*_0_, and previous models (Lande et al. 1994) show that *d*_0_ decreases slowly as the selfing rate increases. Therefore, here I assume *d*_0_ is the same for populations with different selfing rates, and I only consider the influence of the selfing rate on inbreeding depression contributed by slightly deleterious mutations. I denote the haploid genomic mutation rate as *U*, and assume all deleterious mutations are identical with selection coefficient *s* and dominance *h*_*d*_. For a population at the equilibrium state, the inbreeding depression can be approximated by (Roze 2015)

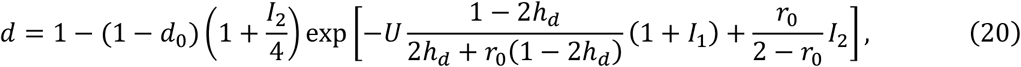

where

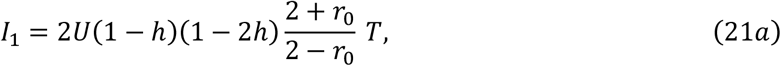

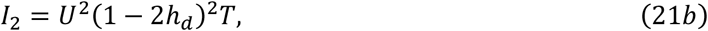

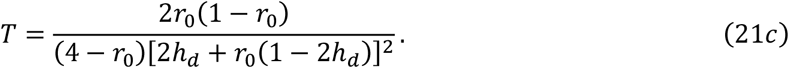

The effect of the background selection on reducing the effective population size (Nordborg et al. 1996) is

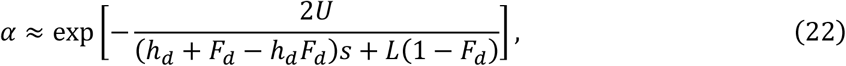

where *L* is the number of crossovers along the chromosomes and *F*_*d*_ = *r*_0_/(2 − *r*_0_). The fixation probability can be calculated by replacing *N*_*e*_ with *αN*_*e*_ in equations (5) and (10).

### Simulations

Simulations are written in R 4.0.2 and the code is archived on Dryad [filed upon acceptance]. Every generation, reproduction occurs first based on the individual selfing rate, followed by selection. The next generation is drawn from a multinomial distribution based on the expected genotype frequencies after selection. For adaptation from standing variation, I run 6*N*_0_ generations to reach the selection-drift-mutation equilibrium (Hermisson and Pennings 2005) with forward and backward mutations, and after pollen limitation happens, the input of mutations stops. For adaptation in a declining population, the population size in the next generation is drawn from a Poisson distribution with the mean being the product of the current population size and the average fitness. The population is considered as having gone to extinction when *N* = 0, and is considered as rescued when the modifier allele goes to fixation or the population size exceeds 10*N*_0_.

