## Supplementary Information for "The genetic basis of adaptation through the evolution of self-fertilization"

PNAS
www.pnas.org

**Supplementary Information for**
### The genetic basis of adaptation through the evolution of self-fertilization

Kuangyi Xu

Kuangyi Xu


**This PDF file includes:**

Supplementary text
Figures S1 to S9
Table S1
SI References

**Supplementary Information Text**

**Derivation of the effective population size and fixation probability at a selfing modifier locus**

Here I aim to derive the fixation probability of a selfing modifier allele in a finite population. To do this, I first obtain the effective population size at a selfing rate modifier locus by deriving the expected allele frequency change and its variance between two discrete generations. By adopting the diffusion method, I then develop an analytical model for the fixation probability and fixation time of a selfing modifier allele, which are compared to results from simulations.

#### 1. Expected allele frequency change and its variance in a finite population

Consider a hermaphrodite population with non-overlapping generations, with population size being $N_t$ at generation $t$. There is a selfing modifier locus with two alleles $A$ and $a$. The frequency of allele $A$ is denoted by $p$, and the frequency of genotype $AA, Aa$ and $aa$ are denoted as $P, 2Q$ and $R$, which can be expressed as $pF+(1-F)p^2$, $2p(1-p)(1-F)$ and $(1-p)F+(1-F)(1-p)^2$. $F$ is the inbreeding coefficient which measures the deviation from Hardy-Weinberg equilibrium. The three genotypes $AA, Aa,$ and $aa$ respectively increase the individual selfing rate by $r, hr$ and 0, where $h$ is the dominance coefficient. I also allow there to be a background selfing rate $r_0$, so the three genotypes $AA, Aa$ and $aa$ will respectively have selfing rate as $r_{AA}=r_0+r, r_{Aa}=r_0+hr$ and $r_{aa}=r_0$.

After reproduction, the offspring population can be divided into two parts based on whether they are produced from selfing or outcrossing. The expected proportion of self-fertilized seeds in the population is $S=Pr_{AA}+2Qr_{Aa}+Rr_{aa}$, and the remaining proportion $T=1-S$ is outcrossed. Among self-fertilized seeds, the frequency of allele $A$ is

$$p_S=\frac{Pr_{AA}+Qr_{Aa}}{S}. \tag{S1}$$

Therefore, the inbreeding coefficient within the self-fertilized seeds $F_S$ is

$$F_S=1-\frac{Qr_{Aa}}{S}\frac{1}{p_S(1-p_S)}\ . \tag{S2}$$

Among seeds produced from outcrossing, the allele frequency should be separately considered for the female (ovule) and male (pollen) gamete pool. The expected frequency of allele $A$ in the female gamete pool $p_T^{(f)}$ is

$$p_T^{(f)}=\frac{P(1-r_{AA})+Q(1-r_{Aa})}{T}. \tag{S3}$$

As selfing event will consume pollen and may result in a decrease in the amount of pollen exported for outcrossing, which is called pollen discounting (Harder and Wilson 1998, Porcher and Lande 2005), I assume individuals with a selfing rate $r$ export a proportion of $1-cr$ relative to those that are completely outcrossing, where $c$ measures the degree of pollen discounting. Therefore, the frequency of allele $A$ in the pollen pool $p_T^{(m)}$ is

$$p_T^{(m)}=\frac{P(1-cr_{AA})+Q(1-cr_{Aa})}{P(1-cr_{AA})+2Q(1-cr_{Aa})+R(1-cr_{aa})}\ . \tag{S4}$$

I assume population is randomly pollinated, so the expected allele frequency in the outcrossed offspring is merely the average of that among the female and male gamete pool as $p_T=\left(p_T^{(f)}+p_T^{(m)}\right)/2$.

Another important genetic factor that constraints the evolution of selfing is inbreeding depression. Inbreeding depression results from the fact that selfing will expose more deleterious recessive mutations to be homozygous than will outcrossing do, thus selfed offspring will on average have lower fitness than outcrossed offspring. In the population at generation $t+1$, the actual proportion of offspring produced from selfing is denoted as $l$, and its expectation is

$$\lambda = E[l] = \frac{(1-d)S}{(1-d)S+T}, \tag{S5}$$

where $d$ incorporates inbreeding depression (Charlesworth and Willis 2009), measured as the fitness decrease of selfed seeds relative to those produced from outcrossing The expected allele frequency change $M_{\Delta p}$ and its variance $V_{\Delta p}$ after one generation are

$$M_{\Delta p} = E[p' - p] = E\big[E[p'(l) - p|l]\big] = \lambda p_S + (1-\lambda)p_T - p, \tag{S6a}$$

$$V_{\Delta p} = Var(p' - p) = E[Var(p'(l) - p|l)] + Var(E[p'(l) - p|l]), \tag{S6b}$$

where $E[\cdot\ |l]$ and $Var(\cdot\ |l)$ are the expectation and variance given that the proportion of selfed offspring is $l$. It can be seen from Eq. (S6a) that the expectation of allele frequency change $M_{\Delta p}$ has been analytically derived, and we only need to focus on the variance $V_{\Delta p}$.

In Eq. (S6b), the first term stands for the variance of allele frequency change given that the proportion of selfed offspring is fixed, and the second term accounts for the effects due to variation of the actual proportion of self-fertilized offspring. Since both the allele frequency and inbreeding coefficient differ between selfed and outcrossed seeds, the two terms can be further partitioned as

$$E[Var(p'(l) - p|l)] = E\big[l^2 V_{\Delta p}^{(S)} + (1-l)^2 V_{\Delta p}^{(T)}\big], \tag{S7a}$$

$$Var(E[p'(l) - p|l]) = Var(lp_S + (1-l)p_T) = (p_S - p_T)^2 Var(l), \tag{S7b}$$

where the $V_{\Delta p}^{(S)}$ and $V_{\Delta p}^{(T)}$ are variance of allele frequency change among selfed and outcrossed offspring.

Before analyzing $V_{\Delta p}^{(S)}$ and $V_{\Delta p}^{(T)}$, recall the key result in Kimura and Crow (1963): when the offspring number of each individual in the population is identically distributed, the variance of allele frequency change after one generation from $t$ to $t+1$ is

$$Var(p' - p) = \frac{p(1-p)}{2N_t\bar{k}}\left[\frac{N_t}{N_t - 1}\frac{V_k}{\bar{k}}(1+F) + (1-F)\right], \tag{S8}$$

where $\bar{k}$ is the average gamete number that each individual contributes to the next generation, and $V_k$ is its variance. Applying the result, we can get the variance of allele frequency change among selfed offspring $V_{\Delta p}^{(S)}$. Given that the proportion of selfed offspring in $t+1$ generation is $l$, the average gamete number contributed through self-fertilization is $2lk$, where $k = N_{t+1}/N_t$ is the average offspring number per individual. Denote the variance of the gamete number contributed by selfing as $V_k^{(S)}$, based on Eq. (S8), $V_{\Delta p}^{(S)}$ is

$$V_{\Delta p}^{(S)} = \frac{p_S(1-p_S)}{2N_t 2kl}\left[\frac{N_t}{N_t - 1}\frac{V_k^{(S)}}{2kl}(1+F_S) + (1-F_S)\right]. \tag{S9}$$

For the variance of allele frequency change contributed through outcrossing $V_{\Delta p}^{(T)}$, it can be decomposed as

$$V_{\Delta p}^{(T)} = Var\left(\frac{p_T^{(f)} + p_T^{(m)}}{2} - p\right) = \frac{V_{\Delta p}^{(f)} + V_{\Delta p}^{(m)}}{4}, \tag{S10}$$

where $V_{\Delta p}^{(f)}$ and $V_{\Delta p}^{(m)}$ are variance of allele frequency change in female and male gametes that participate into outcrossing reproduction. Since both female and male parents contribute an average of $(1-l)k$ gamete into the offspring population, according to Eq. (S8), we have

$$V_{\Delta p}^{(f)} = \frac{p_T^{(f)}\left(1 - p_T^{(f)}\right)}{2N_t k(1-l)}\left[\frac{N_t}{N_t - 1}\frac{V_k^{(f)}}{k(1-l)}\left(1 + F_T^{(f)}\right) + \left(1 - F_T^{(f)}\right)\right], \tag{S11a}$$

$$V_{\Delta p}^{(m)} = \frac{p_T^{(m)}\left(1 - p_T^{(m)}\right)}{2N_t k(1-l)}\left[\frac{N_t}{N_t - 1}\frac{V_k^{(m)}}{k(1-l)}\left(1 + F_T^{(m)}\right) + \left(1 - F_T^{(m)}\right)\right], \tag{S11b}$$

where $V_k^{(f)}$ and $V_k^{(m)}$ are variance of gamete number contributed by each female and male individual (note that although individuals are hermaphroditic, they function as females or males during outcrossing). $F_T^{(f)}$ and $F_T^{(m)}$ are inbreeding coefficient within the female and male parent populations, which can be calculated as

$$F_T^{(f)} = 1 - \frac{Q(1-r_2)}{T}\frac{1}{p_T^{(f)}\left(1-p_T^{(f)}\right)}, \tag{S12a}$$

$$F_T^{(m)} = 1 - \frac{Q(1-\pi r_2)}{P(1-\pi r_1)+2Q(1-\pi r_2)+R(1-\pi r_3)}\frac{1}{p_T^{(m)}\left(1-p_T^{(m)}\right)}. \tag{S12b}$$

Until now, all the components of the overall variance $V_{\Delta p}$ have been presented. However, the expression is quite complex and depends on both the distribution of the gamete number and proportion of self-fertilized offspring $l$.

For a special case when each individual produces a large number of seeds, so that the selfed and outcrossed seeds competing with each other to survive to the next generation, the proportion of selfed offspring in offspring population is subject to a binomial distribution $B(N_{t+1}, \lambda)$. Furthermore, if the offspring number of both selfing and outcrossing reproduction is subject to a Poisson distribution, given a certain value of $l$, the variance of gamete number through selfing is $V_k^{(S)} = 4kl$, and the variance of female and male gamete number participating in outcrossing are $V_k^{(f)} = V_k^{(m)} = k(1-l)$. In this case, substituting the variance into Eq. (S11) and assuming the parent population size $N_t$ not too small, we have

$$V_{\Delta p}^{(S)} = \frac{p_S(1-p_S)}{4N_{t+1}l}(3+F_S), \tag{S13a}$$

$$V_{\Delta p}^{(T)} = \frac{p_T^{(f)}\left(1-p_T^{(f)}\right)+p_T^{(m)}\left(1-p_T^{(m)}\right)}{4N_{t+1}(1-l)}. \tag{S13b}$$

As a result, according to Eq. (S7b), the total variance of allele frequency change after one generation is

$$V_{\Delta p} = \frac{1}{4N}\left[\lambda p_S(1-p_S)(3+F_S) + (1-\lambda)p_T^{(f)}\left(1-p_T^{(f)}\right) + p_T^{(m)}\left(1-p_T^{(m)}\right) + 4\lambda(1-\lambda)(p_S-p_T)^2\right]. \tag{S14}$$

#### 2. Diffusion approximation

I adopt the diffusion limit to solve for the fixation probability and average time to fixation of the selfing modifier allele $A$. Following the method in Crow and Kimura (1970), suppose for a small time interval $\Delta t$, the effects of $AA$ and $Aa$ on individual selfing rate are $r\Delta t$ and $hr\Delta t$. Assume the population size $N$ is constant and the successful gamete number is subject to a Poisson distribution, by taking the limit $\Delta t \to 0$, the infinitesimal change $M$ and variance $V$ can be derived from Eqs. (S6a) and (S14) as

$$M = \lim_{\Delta t \to 0}\frac{M_{\Delta p}}{\Delta t} = Rp(1-p)[h+F-hF+(1-F)(1-2h)\,p], \tag{S15a}$$

$$V = \lim_{\Delta t \to 0} V_{\Delta p} = \frac{p(1-p)}{2N_e}, \tag{S15b}$$

where

$$R = \frac{1-2d-c(1-2dr_0)}{2(1-dr_0)(1-cr_0)}r, \tag{S16a}$$

$$N_e = \frac{2(1-dr_0)}{2+[1-3d+F(1-d)]r_0}N. \tag{S16b}$$

$R$ can be effectively considered as the selection coefficient of the selfing modifier allele $A$, which decreases with the level of pollen discounting and inbreeding depression since

$$\frac{\partial R}{\partial c} = -\frac{1-r_0}{2(1-cr_0)^2(1-dr_0)}r < 0, \tag{S17a}$$

$$\frac{\partial R}{\partial d} = -\frac{2-(1+c)r_0}{2(1-cr_0)(1-dr_0)^2}r < 0. \tag{S17b}$$

Also, Eq.(S16a) shows that when there is no pollen discounting ($c = 0$), the selfing modifier will be favored only when inbreeding depression is lower than 0.5 (Lande and Schemske 1985).

Eq. (S16b) shows that the infinitesimal effective population size does not explicitly depend on pollen discounting $c$, because the third term in Eq. (S14) vanishes during the infinitesimal approximation, although pollen discounting will indirectly affect effective population size by changing $F$. Also, the effective population size is equal to the census size $N$ when the background selfing rate $r_0$ is 0, irrespective of the level of inbreeding depression $\delta$ and inbreeding coefficient $F$.

Based on the above results, the probability of fixation $u(p)$ of the allele $A$ with initial frequency $p$ in a finite population can be calculated as (Kimura 1962)

$$u(p) = \frac{\int_0^p G(x)dx}{\int_0^1 G(x)dx}, \tag{S18}$$

where

$$G(x) = \exp\left(-\int \frac{2M}{V}dx\right). \tag{S19}$$

Conditioned on fixing, the average number of generations until fixation of allele $A$ starting from a frequency $q$ is (Kimura and Ohta 1969)

$$T(p) = \frac{1-u(p)}{u(p)}\int_0^p \psi(x)u(x)dx + \int_p^1 \psi(x)\big(1-u(x)\big)dx\,, \tag{S20}$$

where

$$\psi(x) = \frac{4N_e\int_0^1 G(z)dz}{x(1-x)G(x)}. \tag{S21}$$

**3. Time-separation and approximate solutions**

From Eqs. (S15) and (S16b), we see that both the infinitesimal change and effective population size depend on the inbreeding coefficient $F$. However, $F$ will change as the frequency of the modifier allele changes, so we need to express $F$ as a function of the allele frequency $p$. To do this, we adopt a separation of time scales, which assumes the inbreeding coefficient reaches an equilibrium much faster than does the change of the allele frequency. This assumption has been proved to be effective in previous studies (e.g., Hartfield and Glémin 2016). To allow the inbreeding coefficient to reach an equilibrium, we need to make the allele selectively neutral, so that the allele frequency does not change. For a mating-unrelated locus, this can be done by directly setting the selection coefficient to be 0 (Caballero and Hill 1992). However, this method does not apply to a selfing modifier allele, as the level of inbreeding should change with the allele frequency. Therefore, I choose to adjust inbreeding depression to make the modifier allele selectively neutral.

Here I give a brief description of the derivation of $F$. Note that the selfing rate of genotype $AA$, $Aa$ and $aa$ are $r_0 + r$, $r_0 + hr$ and $r_0$, we can set a virtual inbreeding depression $d_v$ for the extra selfing rate contributed by allele $A$, while the inbreeding depression of selfed offspring due to the background selfing rate is still $d$. In other word, among the selfed offspring of genotype $AA$, a proportion $r$ has fitness $1 - d_v$, while the remaining proportion $r_0$ has fitness $1 - d$. Similarly, the fitness of selfed offspring from genotype $Aa$ can be partitioned as $(1 - d_v)hr$ and $(1 - d)r_0$. By solving the recursion of allele frequency in Eq. (S6a), the virtual inbreeding depression $d_v$ that makes the allele $A$ selective neutral is

$$d_v = \frac{1}{2}\frac{1-c}{1-c(2hpr+(1-2h)Pr+r_0)}, \tag{S22}$$

where $P = pF + (1 - F)p^2$ is the frequency of genotype $AA$. Therefore, the equilibrium inbreeding coefficient $F$ can be solved by substituting $d_v$ into recursions of genotype frequency, which is a complex function of allele frequency $p$.

The dependency of $F$ on allele frequency $p$ makes it difficult to derive an analytical expression for the fixation probability. Therefore, ignoring terms involving $p$ and the higher order term $o(r^2)$, we can further approximate $F$ as

$$F \approx F_0 + \left(2 - \frac{1-c}{1-cr_0}\right)\frac{(1-d)r_0 + h(1-r_0)}{\alpha^2}r, \qquad (S23)$$

where

$$F_0 = \frac{(1-d)r_0}{2-(1+d)r_0}, \qquad (S24a)$$

$$\alpha = 2-(1+d)r_0. \qquad (S24b)$$

$F_0$ is the equilibrium inbreeding coefficient caused by the background selfing rate $r_0$ after accounting for inbreeding depression.

#### 4. Validity of the model

To verify the validity of our model, we compare predictions of the fixation probability from Eq. (S18) with results from simulations. Fig. S8 shows this model works well under various conditions, which may even apply when $r$ is large. Fig. S9 shows that the validity of the model is robust to the initial allele frequency and initial inbreeding coefficient. Also, Table S1 shows that the approximation of a frequency-independent inbreeding coefficient in Eq. (S23) gives satisfactory predictions.

#### 5. Effects of genetic and ecological factors

We can further examine the effects of inbreeding depression and pollen discounting on the effective population size and fixation probability. Based on the approximation of inbreeding depression $F$ in Eq. (S23), we have

$$\frac{\partial F}{\partial c} = \frac{(1-d)r_0 + h(1-r_0)}{\alpha^2}\frac{1-r_0}{(1-cr_0)^2}r > 0, \qquad (S25a)$$

$$\frac{\partial F}{\partial d} = -\frac{2r_0(1-r_0)}{\alpha^2} - r_0\left(2-\frac{1-c}{1-cr_0}\right)\frac{2[(1-h)+(1+h)r_0] + r_0(1-d)}{\alpha^3}r < 0. \qquad (S25b)$$

Eq. (S25a) shows that a more severe pollen limitation will increase the inbreeding coefficient, which is easy to understand as higher pollen discounting makes the selfing enhancer allele $A$ less likely to participate in outcrossing.

Based on the expression of $N_e$ in Eq. (S16b), we have

$$\frac{\partial N_e}{\partial c} \propto -(1-d)(1-dr_0)r_0\frac{\partial F}{\partial c} < 0, \qquad (S26a)$$

$$\frac{\partial N_e}{\partial d} \propto r_0(1-r_0)(1+F)\frac{\partial F}{\partial d} < 0. \qquad (S26b)$$

Eq. (S26a) shows that a stronger pollen discounting decreases the effective population size, since it increases the level of inbreeding. On the other hand, although a higher inbreeding depression will lower the actual level of inbreeding, which increases the effective population size of a mating-unrelated locus, it actually decreases the effective population size at a selfing modifier locus, as shown in Eq. (S26b).

Making use of the approximate solution for the fixation probability $u(p) = 4N_eRh_ep$, it can be seen that

$$\frac{\partial u}{\partial c} = \frac{\partial N_e}{\partial c}Rh_e + \frac{\partial R}{\partial c}N_eh_e + N_eR\frac{\partial h_e}{\partial c} < 0, \qquad (S27a)$$

$$\frac{\partial u}{\partial d} = \frac{\partial N_e}{\partial d}Rh_e + \frac{\partial R}{\partial d}N_eh_e + N_eR\frac{\partial h_e}{\partial d} < 0. \qquad (S27b)$$

As expected, more severe inbreeding depression or pollen discounting will make the selfing modifier less advantageous, thus lowering its fixation probability.

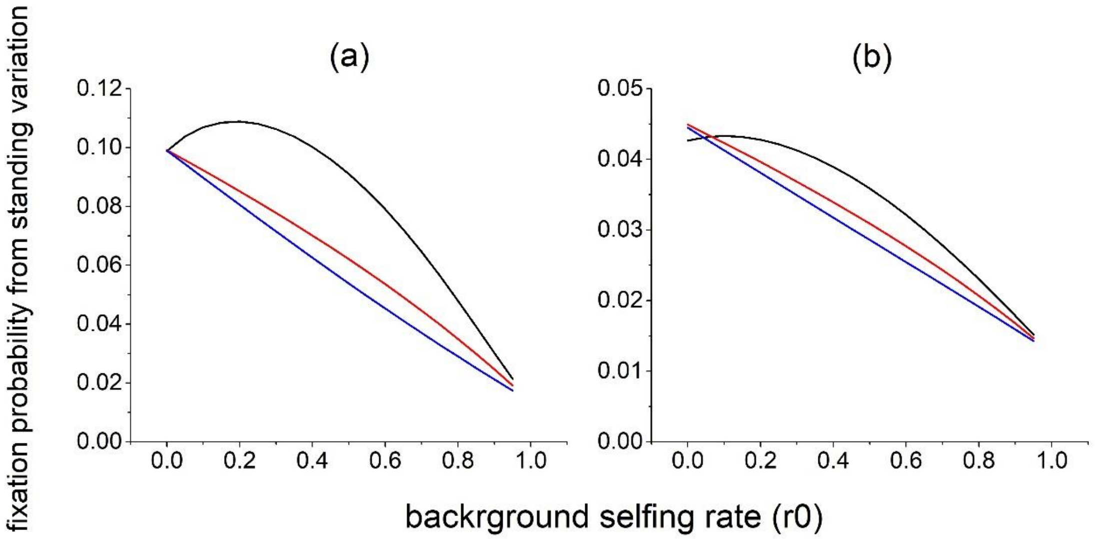


**Fig. S1.** The effects of different factors on the fixation probability from standing variation. Parameters are: panel (a), $d = 0.8$; panel (b), $N = 5000$. Unless specified, parameters are the same as those in Fig. 2(b). Recessive alleles have more advantages in fixation when $\boldsymbol{d}$ is larger (compare Figs. S1(b) and 2(a)), and population size is larger (compare Figs. S1(a) and 2(b)).

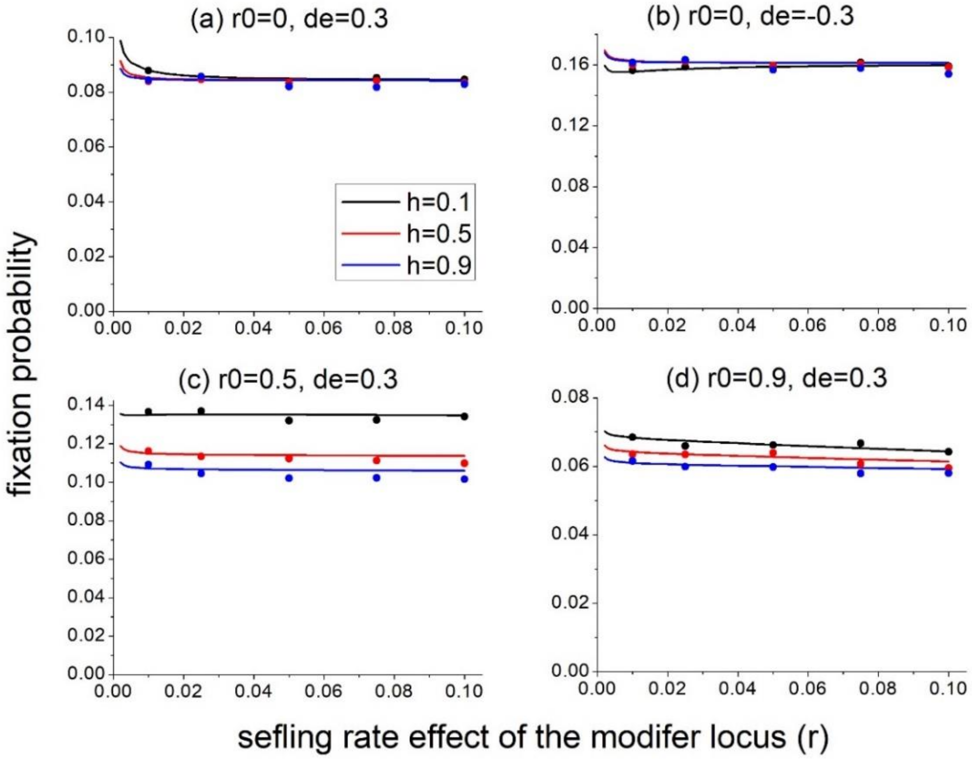


**Fig. S2.** Influences of selfing rate effects of the modifier locus on fixation probability from standing variation. Generally, fixation probability is nearly independent of $\boldsymbol{r}$. Lines are model predictions from equation (10), and dots are simulation results. For all panels, $r = 0.01, d = 0.6, N = 20000$, $\nu_1 = \nu_2 = 10^{-6}$.

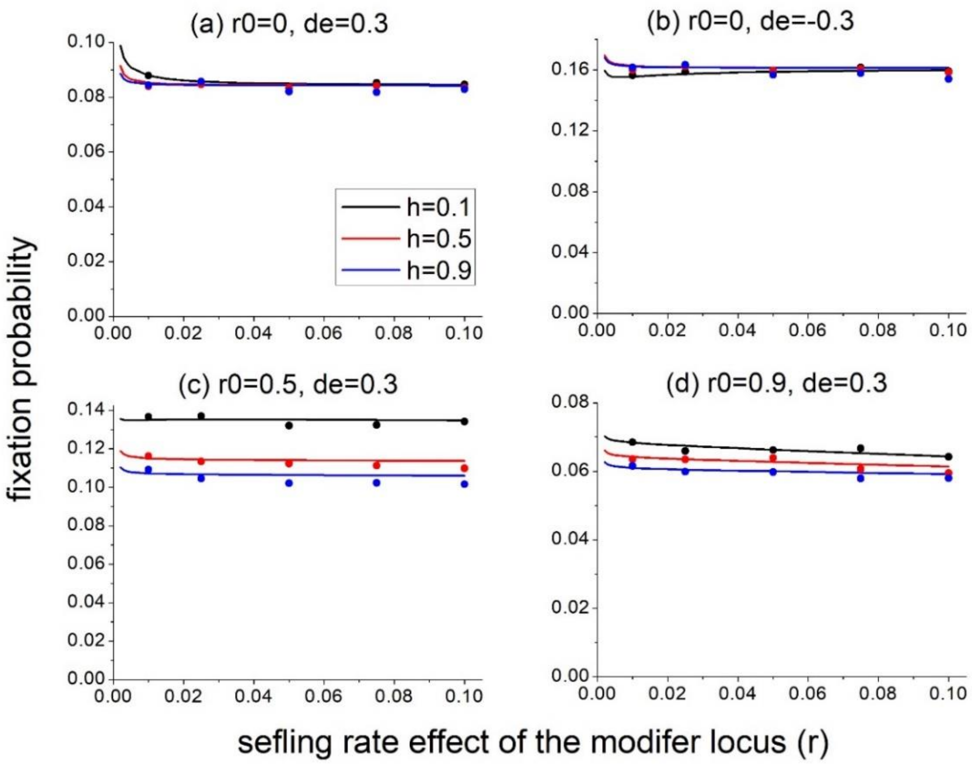


**Fig. S2.** Influences of selfing rate effects of the modifier locus on fixation probability from standing variation. Generally, fixation probability is nearly independent of $\boldsymbol{r}$. Lines are model predictions from equation (10), and dots are simulation results. For all panels, $r = 0.01, d = 0.6, N = 20000$, $\nu_1 = \nu_2 = 10^{-6}$.

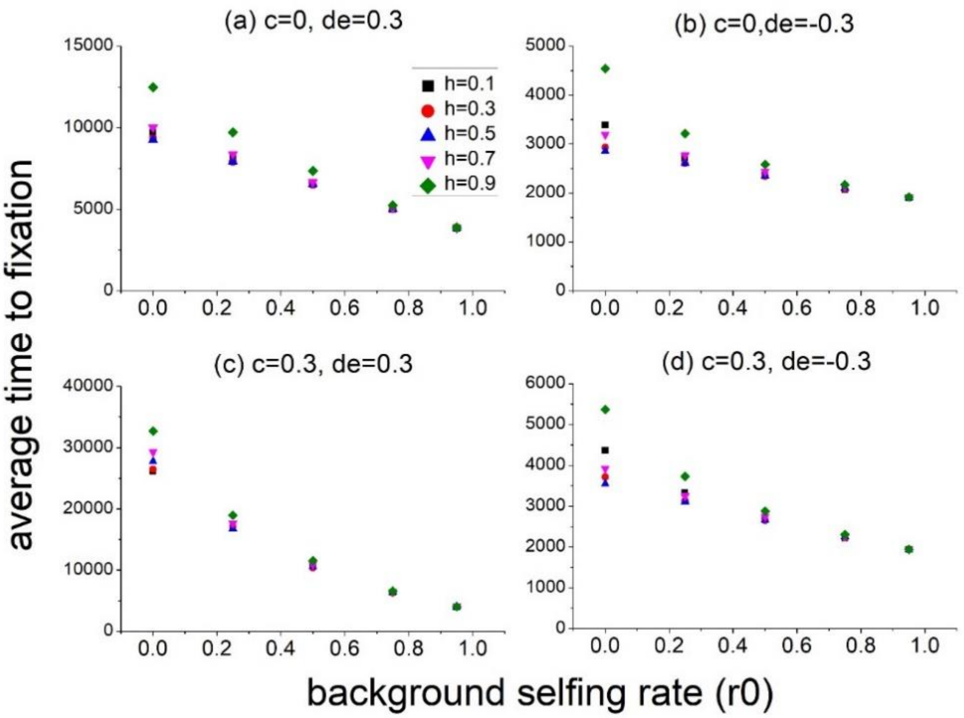


**Fig. S3.** Average time towards fixation from standing variation in a constant population. Results are from simulations. For all panels, $r = 0.01, d = 0.6, N = 20000, \nu_1 = \nu_2 = 10^{-6}$.

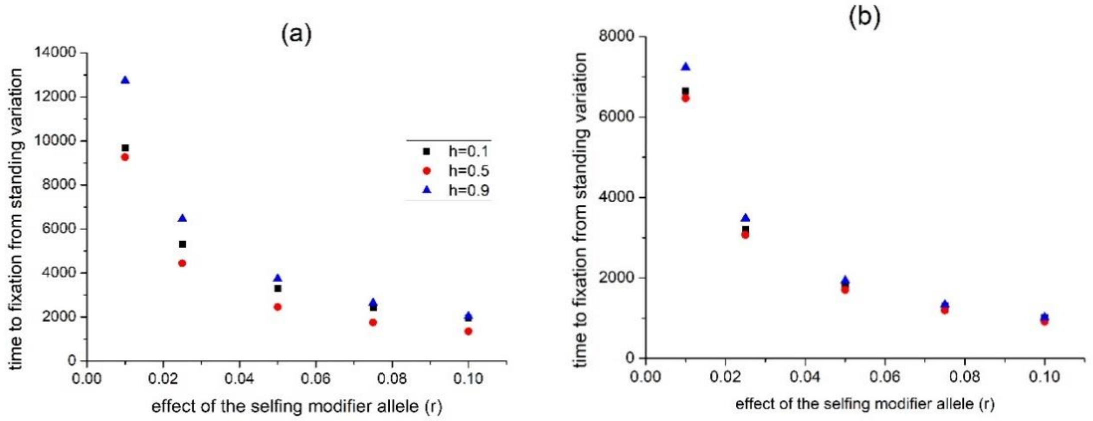


**Fig. S4.** Influences of the effect of the selfing modifier allele $\boldsymbol{r}$ on the average time towards fixation from standing variation. Results are from simulations. $r_0 = 0$ and 0.5 for panels (a) and (b). For both panels, $d = 0.6, d_e = 0.3, N = 20000$. The conclusion that fixation is slower for more dominant alleles holds for different values of $r$.

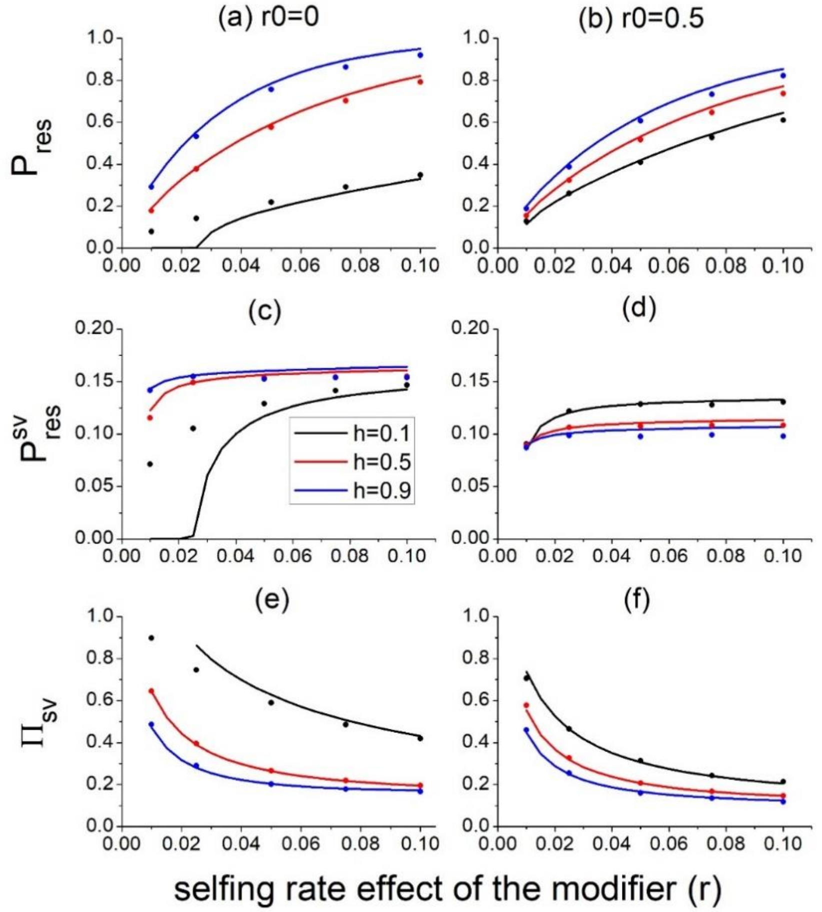


**Fig. S5.** Effects of selfing rate effects of modifier allele on the overall survival probability ($P_{res}$), survival probability from standing variation ($P_{res}^{sv}$), and the relative contribution from standing variation ($\Pi_{sv}$) under demographic decline. Fig. 3(c) and 3(d) show that a small $r$ will favor adaptation through dominant alleles. $r_0 = 0$ for panels (a), (c), (e), and $r_0 = 0.5$ for panels (b), (d), (f). For all panels, $0.6, d_e = -0.3, N = 20000, \lambda = 0.002$.

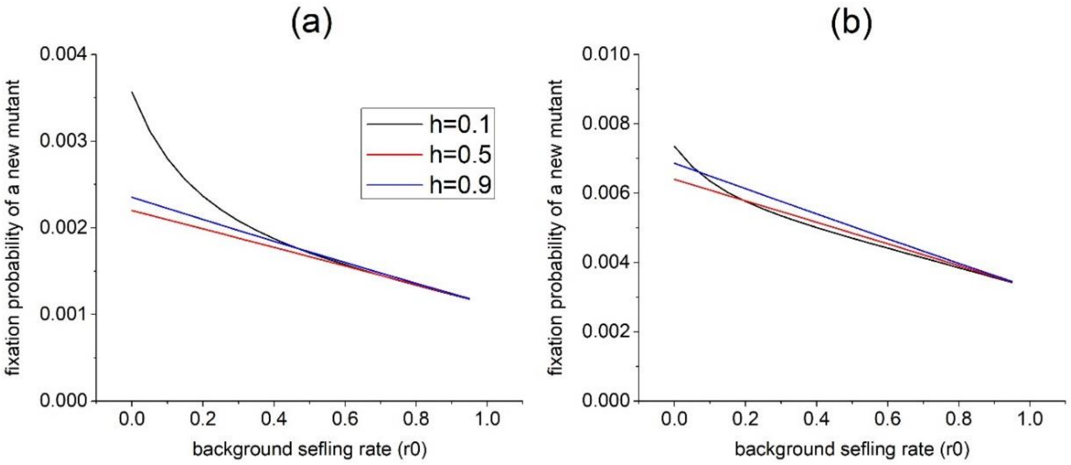


**Fig. S6.** Effects of the background selfing rate on fixation probability from standing variation in a population suffering a bottleneck for mating-unrelated adaptive alleles. The adaptive allele is previously selected against with selection coefficient $s_1$, and later becomes adaptive with selection coefficient $s_2$. Panel (a), $s_1 = -0.01, s_2 = 0.01$; panel (b), $s_1 = -0.01, s_2 = 0.03$. For both panels, $N_0 = 20000, N_c = 500$.

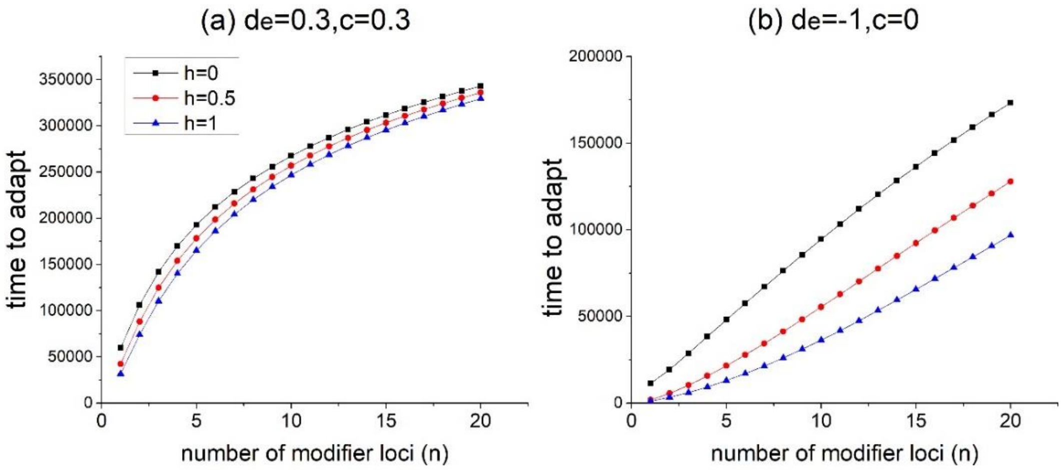


**Fig. S7.** Effects of genetic architecture on the rate of adaptation through new mutations in a constant population. Time to adapt is the time for all modifiers to fix, estimated by equation (19). Panels (a) and (b) show the case of weak and strong selection on selfing modifiers, respectively. The overall increase of the selfing rate is $\Delta r = 0.2$, and $r_0 = 0, N = 100, v_1 = 10^{-5}$.

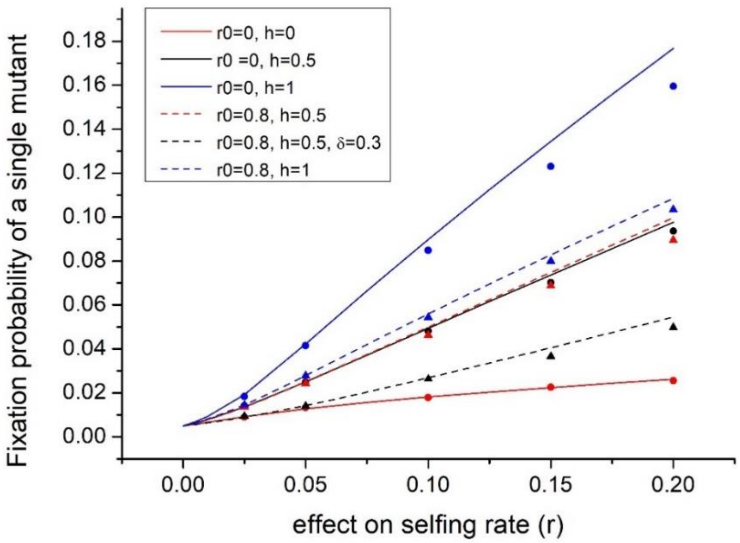


**Fig. S8.** Changes of the fixation probability of a single modifier allele mutant with effect $r$ under different situations. Lines are model predictions from Eq. (S18). The round dots and triangles respectively denote results from simulations under $r_0 = 0$ and $r_0 = 0.8$. Unless specified, $N = 100, c = 0, d = 0$.

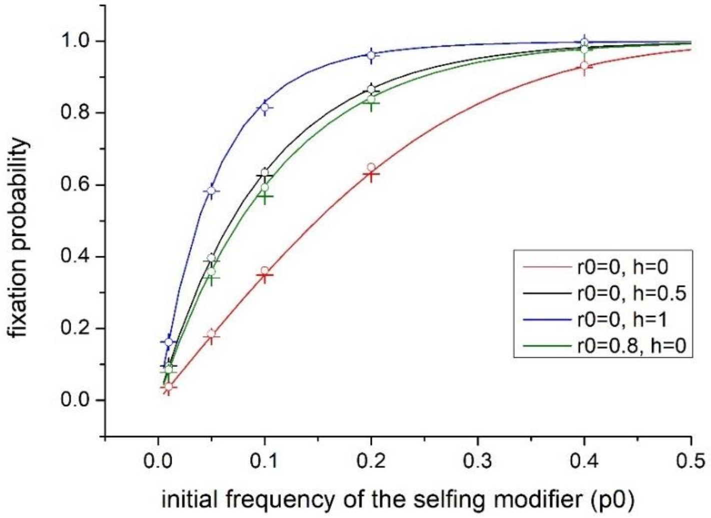


**Fig. S9.** The effects of the initial frequency $p_0$ and initial inbreeding coefficient $F_0$ on the fixation probability. Lines are from model predictions by solving Eq.(S18). For the simulation results, the symbol $+$ denotes $F_0 = 0$, and the open circle denotes $F_0 = 1$. Other parameters used are $r = 0.1, c = 0, d = 0$.

**Table S1.** Comparison of the fixation probability a single mutant allele between accurate solution and approximation. Results shown are scaled as $u(1/2N)\,N$, where $N = 100$. "Exact" is the numeric solution of Eq. (S18), in which $F$ is a function of $p$. "Rel. Err" is the relative error between results given by the frequency-independent approximation in Eq. (S23) to the exact model predictions.

| | $r0 = 0, d = 0, c = 0$ | | | $r0 = 0.8, d = 0, c = 0$ | | | $r0 = 0.8, d = 0, c = 0.3$ | | | $r0 = 0.8, d = 0.3, c = 0$ | | |
|---|---|---|---|---|---|---|---|---|---|---|---|---|
| $r$ | $h = 0$ | $h = 0.5$ | $h = 1$ | $h = 0$ | $h = 0.5$ | $h = 1$ | $h = 0$ | $h = 0.5$ | $h = 1$ | $h = 0$ | $h = 0.5$ | $h = 1$ |
| 0.005 | | | | | | | | | | | | |
| Exact | 0.58 | 0.63 | 0.69 | 0.62 | 0.63 | 0.65 | 0.61 | 0.62 | 0.63 | 0.56 | 0.57 | 0.57 |
| Rel. Err. | -1.2E-4 | -2.7E-7 | 7.3E-5 | -1.0E-5 | -7.4E-7 | 3.6E-6 | -1.7E-6 | 1.8E-5 | 1.5E-5 | -1.1E-5 | -7.0E-7 | 4.5E-6 |
| 0.02 | | | | | | | | | | | | |
| Exact | 0.84 | 1.15 | 1.55 | 1.09 | 1.15 | 1.22 | 1.03 | 1.09 | 1.15 | 0.77 | 0.81 | 0.84 |
| Rel. Err. | -1.5E-3 | -1.3E-5 | 7.0E-4 | -1.9E-4 | -3.9E-5 | 2.5E-5 | -9.2E-5 | 1.9E-4 | 1.3E-4 | -2.1E-4 | -3.9E-5 | 4.4E-5 |
| 0.1 | | | | | | | | | | | | |
| Exact | 1.81 | 4.94 | 8.99 | 4.38 | 4.99 | 5.59 | 4.07 | 4.61 | 5.15 | 2.34 | 2.69 | 3.04 |
| Rel. Err. | -1.5E-2 | -4.7E-4 | 3.4E-4 | -4.9E-3 | -1.4E-3 | 3.9E-4 | -5.1E-3 | 2.6E-4 | 4.1E-4 | -7.0E-3 | -2.2E-3 | 3.0E-4 |
